# Local clocks within human tissues reveal widespread 24-hour rhythms in gene expression

**DOI:** 10.64898/2026.03.24.713830

**Authors:** Teele Palumaa, Jakob M. Cherry, Priit Palta, Angus C. Burns

## Abstract

Circadian clocks govern the 24-hour rhythmic activity of organs across the human body and these nycthemeral rhythms are critical for human physical and mental health. Here, we analysed 14,886 samples across 45 human tissues and inferred tissue-specific local phases using CHIRAL to define the rhythmic transcriptome. Local phase inference identified a median of 5,747 rhythmic transcripts across tissues, including 5,531 across brain tissues, indicating that prior donor-level approaches have substantially underestimated the extent of rhythmic transcription in the human brain. In the brain, rhythmic genes were enriched for neurotransmitter pathways, the synaptic vesicle cycle, circadian entrainment and major neurodegenerative disorders. Strikingly, more than 200 genes in Alzheimer’s, Parkinson’s, Huntington’s and prion disease pathways displayed significant 24-hour rhythmicity across cortical, striatal and hippocampal regions, segregating into distinct diurnal and nocturnal phase clusters. These included Parkinson’s disease genes *SNCA* and *PRKN*, Alzheimer’s disease genes *PSEN1* and *APP*, prion disease genes *PRNP* and *EIF2AK3* (*PERK*), and Huntington’s disease genes *HTT* and *GPR52*. Among these disease-associated rhythmic transcripts were several targets already under therapeutic investigation for which administration at specific times of day may improve therapeutic efficacy or tolerability. Here, we observe that human tissues, including the brain, exhibit distinct local rhythmic organisation. These findings provide novel insights into disease mechanisms and highlight opportunities for target discovery, drug development and chronotherapeutic intervention.

## Introduction

The mammalian circadian system is hierarchical, with the hypothalamic suprachiasmatic nuclei (SCN) acting as the central pacemaker that coordinates a distributed network of self-sustained peripheral oscillators in tissues throughout the human body, each with an intrinsic periodicity of ∼24 hours [1, 2]. The primary zeitgeber that entrains the central pacemaker is light exposure [3, 4]. However, peripheral oscillators retain tissue-specific dynamics and are capable of being entrained or masked by external periodic factors such as exercise, feeding, sleep, glucocorticoids, and temperature, such that they need not be phase-locked to the central pacemaker [5-8].

These tissue-specific clocks shape diverse physiological processes ranging from metabolism and immune function to synaptic plasticity and neuroendocrine signalling [9-12], and the disruption of circadian rhythms is associated with cardiometabolic diseases, neurodegenerative diseases, sleep and psychiatric disorders [13-18]. Many of the most commonly prescribed drugs target proteins that are known to be rhythmic in mice and chronotherapeutic approaches have shown promise in several areas of medicine, including cardiology and oncology [19-21]. Given their importance for human disease, there is a translational need for characterising the rhythmic transcriptome of human tissues.

Earlier efforts to map human rhythmic transcriptomes have established that temporal information can be recovered from unordered samples in a tissue-specific manner to assess rhythmicity in a handful of human tissues, though with limited success in the brain [22, 23]. Complementary studies that anchor expression by time-of-death have provided insight into brain rhythmic transcriptomes, despite impaired precision arising from the variance in phase-angles of entrainment among free-living human beings [24-31]. More recently, Talamanca, Gobet [32] introduced CHIRAL, a method that precisely estimates phase by leveraging a compact set of validated circadian clock transcripts. The authors estimated a single donor-level global phase to assess rhythmicity across tissues in the GTEx database. A key conclusion from this approach was that peripheral metabolic tissues exhibit the strongest rhythmic transcriptomes, whereas brain tissues showed weaker rhythms in both the number of rhythmic transcripts and their amplitudes. However, as yet this method has not been applied at the level of individual tissues.

In this study, we estimated local phase for individual human tissues in GTEx v11 and assessed 24-hour rhythms in the human transcriptome. We observed a median of 5,747 rhythmic transcripts across tissues, with a median of 5,531 rhythmic transcripts in brain tissues. We present findings with significant implications for disease mechanisms and drug development in neurodegenerative diseases.

## Results

We leveraged 14,886 RNA-seq samples across 45 human tissues from 927 donors in GTEx v11. We estimated the local phase (LP) for each tissue sample from each donor, and the global phase (GP) for each donor, using the circular hierarchical reconstruction algorithm and a set of 12 conserved clock genes (CHIRAL) [32]. Across tissues, including the brain, the estimated LPs were systematically phase-shifted relative to GP, and the strength of alignment was low to moderate (*r* range = 0.20-0.70, median *r* = 0.50, Fig. 1A). Using LPs, we detected a median of 5,747 rhythmic transcripts across tissues, including a median of 5,531 rhythmic transcripts in brain tissues (*P*_*Bonferroni*_ < 0.05; Fig. 1B). We found that rhythmic genes included both protein-coding and non-coding transcripts (17,240 and 8,985 transcripts, respectively), with 4,379 genes rhythmically expressed across ≥20 tissues; notably, seven genes were rhythmic in every tissue, all of which are key components of the molecular circadian clock (*BMAL1, CIART, CRY2, DBP, PER1, PER3*, and *TEF)* (Fig. 1C-D). The striatal orphan G-protein-coupled receptor *GPR52* displayed selective rhythmicity in the nucleus accumbens (Fig. 1E). *GPR52* is known to modulate huntingtin (*HTT*) levels in the striatum and is a current drug target for treating Huntington’s disease [33-35]. Remarkably, the gene for huntingtin (*HTT*) exhibited phase-locked rhythmic expression in the nucleus accumbens with *GPR52* (*GPR52* φ = 1.57 h, *HTT* φ *=* 1.42 h; Fig. 1E).

**Figure 1.**
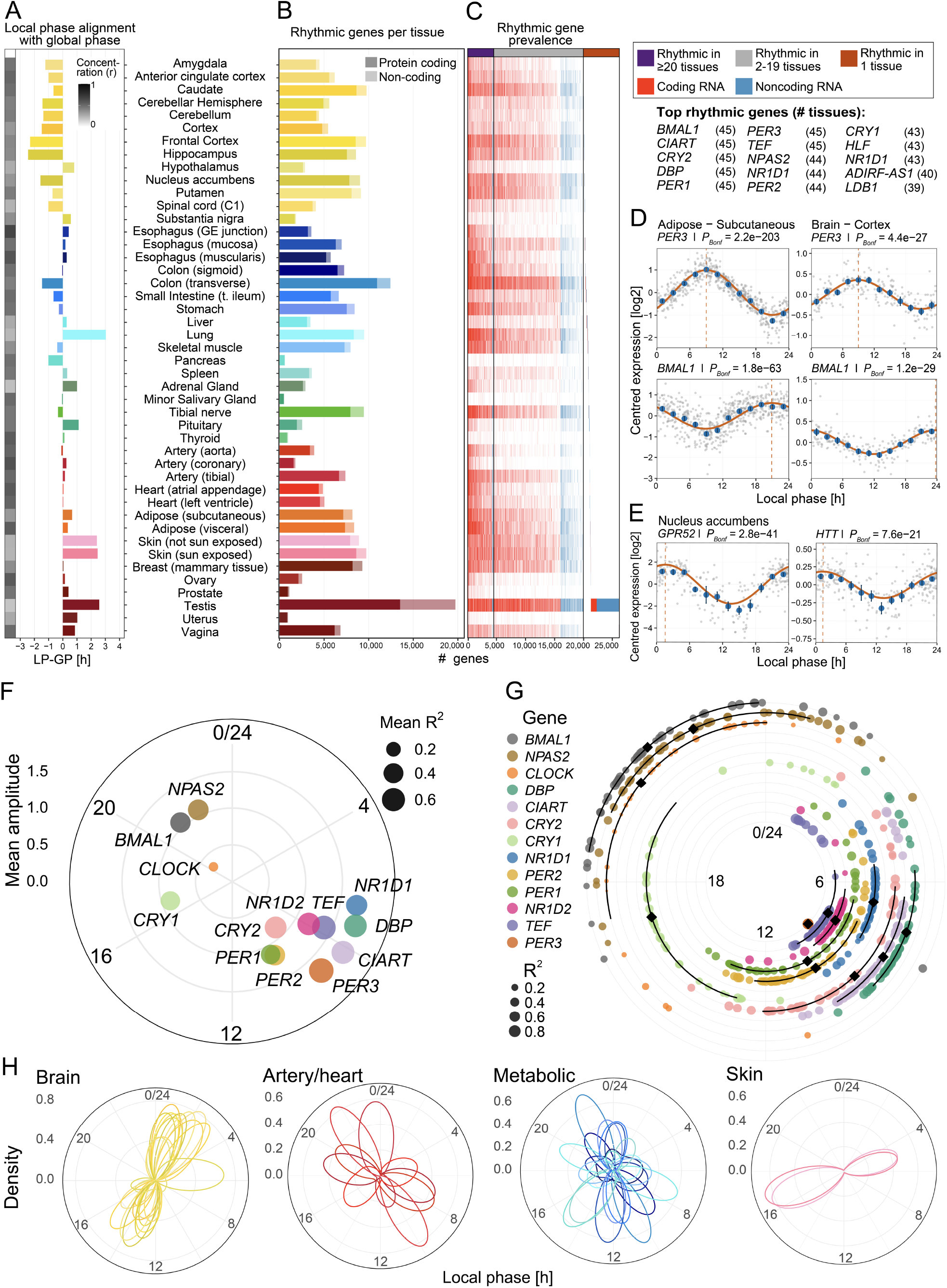
Widespread and tissue-specific circadian gene expression across human tissues. (A) Difference between tissue local phase (LP) and donor global phase (GP) across tissues, shown as the mean LP-GP offset in hours, with grayscale colour indicating phase concentration (*r*). (B) Number of rhythmic genes per tissue, stratified into protein-coding and non-coding genes. (C) Prevalence of rhythmic genes across tissues, grouped by genes rhythmic in 1 tissue, 2–19 tissues, or ≥ 20 tissues, and annotated by protein-coding versus non-coding biotype. The top 15 rhythmic genes are highlighted with the number of tissues in which they were rhythmic. (D, E) Harmonic regression fits for *PER3* and *BMAL1* in subcutaneous adipose tissue and cerebral cortex (D), and *GPR52* and *HTT* in the nucleus accumbens (E). (F) Mean phase and amplitude of canonical clock genes across tissues in which they were rhythmic, with point angle indicating the mean phase. (G) Distributions of rhythmic clock gene phases across tissues, with each point representing a rhythmic gene in a tissue; black symbols indicate the circular mean, and black arcs the ± SD. (H) Examples of tissue-group phase density distributions, illustrating the distribution of rhythmic gene phases across selected organ systems. In (D, E), grey points show individual samples; blue points and error bars show the mean ±SEM within 2-h bins; curves show the fitted harmonic regression model; vertical dashed lines indicate the estimated peak phase.

Across tissues, core clock genes displayed a highly structured phase distribution, with the positive-arm components of the circadian clock transcriptional-translational feedback loop, *BMAL1, CLOCK*, and *NPAS2*, clustering in the late night to early morning and *PER, CRY, NR1D, DBP, TEF* and *CIART* genes occupying the opposing phase domain, consistent with the conserved phase-ordering of clock genes from diurnal mammals [36] (Fig. 1F-G). The phase distributions of rhythmic genes varied across tissues but generally showed bimodal clustering with day and night modules. Brain tissues showed coherent cross-tissue bimodal clustering at midday and midnight, whereas cardiovascular and metabolic tissues exhibited broader phase organisation with peaks in the morning and evening, and skin tissues displayed a pattern centred in the early morning and late afternoon (Fig. 1H).

The most commonly enriched pathway for rhythmic genes across the 45 tissues was the circadian rhythms pathway (Fig. 2A). Among the top 20 most commonly enriched pathways across all tissues were genes involved in endocytosis, protein processing in the endoplasmic reticulum, sphingolipid signalling, autophagy, insulin signalling, bacterial invasion, as well as several cancer-related pathways. In the brain specifically, either or both the circadian entrainment and core circadian rhythms pathways were enriched in all tissues (Fig. 2B). Rhythmic genes were also enriched for major neurotransmitter systems, including dopaminergic, glutamatergic and GABAergic signalling, as well as long-term potentiation and axon guidance. Circadian-related transcription factor pathways were likewise enriched, including genes targeted by CLOCK, LHX1, HIF-2α, HSF1, AHR, PPARγ, USF1, MEF2A/2D and EPAS1, and these transcription factors were themselves frequently rhythmic across tissues [37-42].

**Figure 2.**
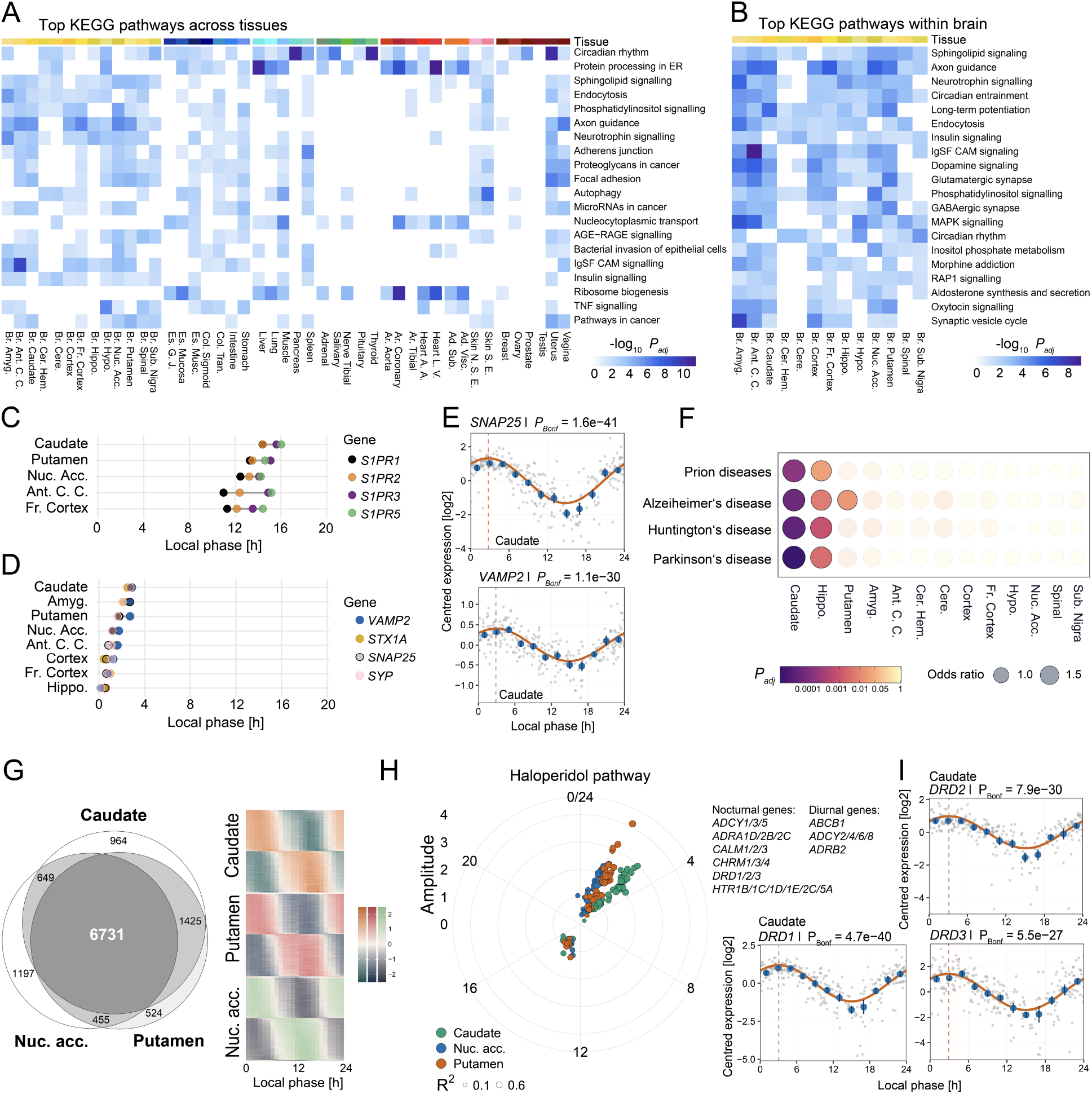
Tissue-wide and brain-specific enrichment of rhythmic pathways. (A, B) Top 20 KEGG pathways enriched among rhythmic genes across all tissues (A), and among rhythmic genes within brain tissues (B). (C, D) Timing of peak rhythmic sphingolipid signalling genes (C) and rhythmic synaptic vesicle cycle genes (D) across brain tissues. (E) Harmonic regression fits for synaptic vesicle cycle genes *SNAP25* and *VAMP2* in the caudate. (F) Enrichment of neurodegeneration-related pathways in brain tissues. (G) Overlap of rhythmic genes among the three striatal tissues: nucleus accumbens, caudate and putamen. The adjacent heatmap shows the centred expression profiles across the local phase for genes rhythmic in all three tissues. (H) Phase-amplitude distribution of genes from the haloperidol signalling pathway that were rhythmic in the nucleus accumbens, caudate and putamen. (I) Harmonic regression fits for dopamine receptor genes from the haloperidol pathway. In (E, I), grey points show individual samples; blue points and error bars show the mean ±SEM within 2-h bins; curves show the fitted harmonic regression model; vertical dashed lines indicate the estimated peak phase.

In twelve of thirteen brain tissues, the sphingolipid signalling pathway was enriched for rhythmic genes. Recent work has established a role for glial sphingolipid signalling in the nycthemeral remodelling of circadian clock neurites, and sphingolipids themselves exhibit diurnal variation in plasma [43, 44]. We identified highly coherent rhythmicity of the sphingosine-1-phosphate receptors (*S1PR*) 1-3 and 5, with each peaking in the day with apparent phase ordering of *S1PR1 & S1PR2* preceding *S1PR3 & S1PR5* (Fig. 2C). The synaptic vesicle cycle pathway was enriched in eight brain tissues and is important in maintaining sleep-wake state stability [45, 46]. We identified a coordinated nocturnal presynaptic vesicle-cycle module in the brain, wherein *VAMP2, SNAP25, STX1A, SYN1* and *SYP* all reached a tightly clustered acrophase in expression between midnight and ∼3am (Fig. 2D & Fig. 2E).

In examining tissue-specific pathway enrichment in the brain, we observed several neurodegenerative disease pathways enriched in striatal and hippocampal brain regions, including Alzheimer’s (AD), Parkinson’s (PD), Huntington’s (HD) and prion diseases (Fig. 2F). Brain-related drug target enrichment identified genes underlying haloperidol action were significantly enriched for rhythmicity, including in the three striatal regions. The caudate, nucleus accumbens and putamen demonstrated a high degree of overlap in rhythmic genes with consistent phases among the shared genes (Fig. 2G). The haloperidol pathway genes formed distinct nocturnal and diurnal clusters with consistent phasing across the striatal tissues (Fig. 2H), with dopamine and serotonin receptor genes peaking in the night and a subset of adenylyl cyclases and the beta-2 adrenergic receptor peaking in the daytime (Fig. 2I).

Given the tissue-specific enrichment of neurodegenerative diseases in the brain, we sought to further probe the rhythmicity of genes underpinning these disorders. First, we examined the rhythmicity of AD and PD genes identified from genome-wide association studies of common and rare variants as well as genes known to cause Mendelian inherited forms of the disorders [47, 48]. We generated “disease clocks” that depict the coherent rhythmicity of these genes across brain tissues in concentric rings (Fig. 3A). The significantly rhythmic AD and PD genes segregated into two clear clusters based on the acrophase of gene expression either in the day or night. For example, the classical Mendelian early-onset AD gene presenilin-1 (*PSEN1*) and *APH1B*, which form critical components of the γ-secretase complex that cleaves amyloid precursor protein (APP), were rhythmic across brain tissues and showed coherent peaks in the daytime, as did *APOE* (Fig. 3B & Fig. 3C) [49, 50]. In contrast, the *APP* transcript itself was rhythmic across brain tissues but reached its acrophase around midnight (mean *APP* φ = 0.88h), consistent with prior work which demonstrated a nycthemeral peak in amyloid beta in the hippocampus and striatum in the night [51].

**Figure 3.**
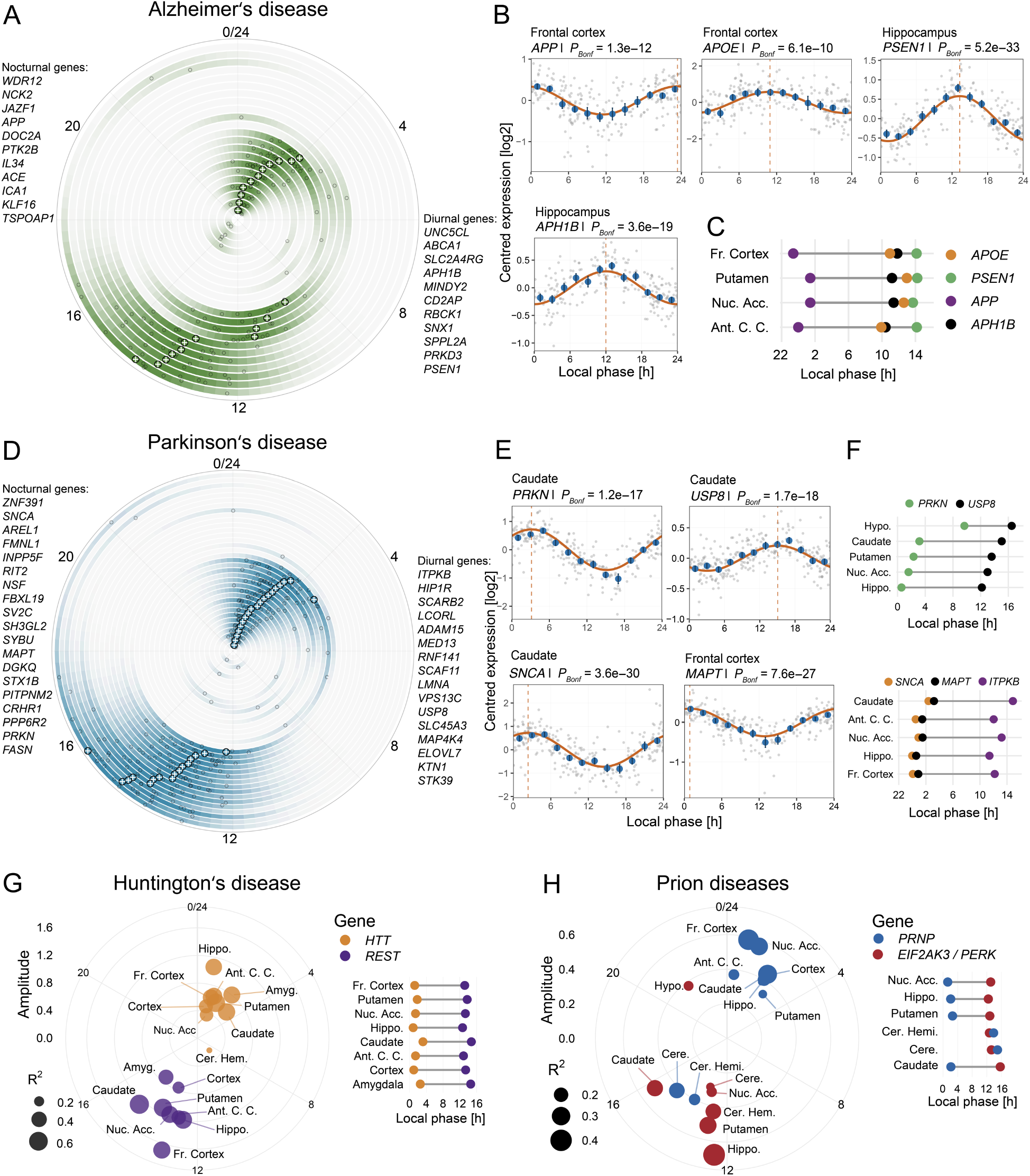
Circadian organisation of neurodegeneration-associated genes across human brain tissues. (A) Alzheimer’s disease “clock” depicting the coherent acrophase distribution of rhythmic genes associated with Alzheimer’s disease across brain tissues. (B) Harmonic regression fits for selected Alzheimer’s disease-associated genes *APP, APOE, PSEN1* and *APH1B* in the frontal cortex and hippocampus. (C) Tissue-specific phase distributions for these genes across brain regions. (D) Parkinson’s disease “clock” depicting the coherent acrophase distribution of rhythmic genes associated with Parkinson’s disease across brain tissues. (E) Harmonic regression fits for selected Parkinson’s disease-associated genes *PRKN, USP8, SNCA* and *MAPT* in caudate and frontal cortex. (F) Tissue-specific phase distributions for these genes and *ITPKB* across brain regions. (G) Phase-amplitude distribution of selected Huntington’s disease-associated genes across brain tissues. The adjacent panel shows tissue-specific phase distributions for *HTT* and *REST*. (H) Phase-amplitude distribution of selected prion disease-associated genes across brain tissues. The adjacent panel shows tissue-specific phase distributions for *PRNP* and *EIF2AK3* (*PERK*). In (A, D), each concentric ring represents one gene, and all brain tissues in which that gene was rhythmic are shown. Small grey points indicate tissue-specific local phases, and white crosses indicate the circular mean phase across tissues. In (B, E), grey points show individual samples; blue points and error bars show the mean ±SEM within 2-h bins; curves show the fitted harmonic regression model; vertical dashed lines indicate the estimated peak phase.

We observed a similar segregation of PD risk genes into diurnal and nocturnal clusters, with similar overall phases of these clusters to their AD counterparts (Fig. 3D). Indeed, the microtubule associated protein tau (*MAPT*) gene, which is involved in the pathogenesis of both AD and PD, was coherently rhythmic in 8 brain tissues with an average peak timing in the night (mean *MAPT* φ = 2.12h; Fig. 3E & Fig. 3F). In terms of PD specific genes, the alpha-synuclein (*SNCA*) gene was coherently rhythmic in 8 brain tissues with a peak in the night (mean *SNCA* φ = 1.02h), whereas *ITPKB*, a component of the inositol pathway which we found to be enriched for brain-rhythmic genes and is implicated in PD via its role in modulating alpha-synuclein, showed a daytime peak (Fig. 3F) [52]. The *PRKN* gene, mutations of which cause an autosomal recessive form of PD, exhibited a coherent nighttime peak in 10 brain tissues [53]. Whereas *USP8*, which promotes the degradation of *PRKN* protein, reached a coherent acrophase across brain tissues during the day (Fig. 3E & Fig. 3F). Together, these findings reveal a temporal architecture of AD and PD risk gene rhythmic expression across the tissues of the human brain, suggesting that neurodegeneration-related pathways are organised into distinct day- and night-peaking programs. These results may be of potential clinical relevance for AD and PD symptom management, vulnerability, mechanism, and chronotherapeutic investigation.

Huntington’s disease and transmissible spongiform encephalopathies (also known as prion diseases) are progressive, fatal neurodegenerative disorders and pathways for these diseases were enriched in brain tissues (Fig. 2F). Sleep-wake disturbance arises preclinically in these diseases and tracks their progression [54, 55]. In addition to the phase-locked rhythmicity of *HTT* with *GPR52* in the nucleus accumbens, *HTT* showed a highly coherent rhythmicity across most brain tissues with acrophases in the night (Fig. 3G). In contrast, the REST transcription factor, which plays a major role in Huntington’s disease pathogenesis, showed rhythmic gene expression antiphase to that of *HTT* and the REST transcription factor target pathway was enriched for rhythmic genes across brain tissues [56, 57]. Similarly, the prion protein gene *PRNP* peaked in the night, whereas *EIF2AK3/PERK*, which is upregulated in disease progression triggering neuronal death, peaked in antiphase to *PRNP* in most tissues (Fig. 3H) [58]. The exception was the cerebellar tissues, which may have relevance to strain-specific neurodegeneration of the cerebellum [54].

## Discussion

Circadian rhythms are a fundamental organising force in physiology, with profound implications for human health, yet the tissue-level architecture of rhythmic gene expression in humans has remained largely inaccessible. Here, we demonstrate that local clocks uncover widespread 24-hour gene expression in the human body. The scale of the effect is striking; by estimating phase at the tissue level, we detect a median of 5,747 rhythmic transcripts across tissues. This reframes the human circadian transcriptome as a locally organised system in which tissue clocks are only partly captured by prior donor-level estimates of phase.

Notably, we observed extensive rhythmic gene expression across brain regions, with enrichment of rhythmic genes in pathways involved in synaptic signalling, axon guidance, the synaptic vesicle cycle, neurotransmission and circadian entrainment. Human brain rhythmicity is difficult to map because direct serial sampling is not possible, yet earlier studies using time of death estimates of phase have suggested that transcripts cycle across cortical and limbic regions and that core clock genes remain phase-coherent outside the suprachiasmatic nucleus [24, 25]. Our results extend that literature substantially: once the local phase of brain tissues was modelled, all 13 showed clear day- and night-peaking programmes and on average five thousand transcripts were rhythmic. These findings contrast with earlier work that attempted to map the rhythmic transcriptome using donor-level phase estimates which suggested that there was more limited rhythmic gene expression in the brain [26, 32]. These findings indicate that the human brain harbours far richer 24-hour transcriptional organisation than previously appreciated.

A particularly striking result was the enrichment of rhythmic genes implicated in major neurodegenerative disorders. A large body of literature has noted rhythmic disturbance in mood, cognition, sleep, and motor symptoms in neurodegenerative disorders such as Parkinson’s disease (PD) and Alzheimer’s disease (AD) that often manifest preclinically and predict the onset of disease [18, 59, 60]. These disturbances are increasingly viewed as bidirectionally linked to neurodegeneration. Our findings provide a molecular framework for this concept by showing that, in the healthy human brain, many genes implicated in neurodegenerative disease are themselves organised into coherent 24-hour programmes across brain tissues, largely represented by nocturnal and diurnal clusters. This temporal structure suggests that disease-relevant processes such as amyloid handling, synuclein homeostasis, mitophagy, neuroimmune signalling and synaptic vesicle cycling may be differentially poised across the day, with potential implications for chronotherapeutic targeting of these pathways.

Indeed, several rhythmic genes highlighted here are already being pursued as therapeutic targets, including *SNCA*/α-synuclein in Parkinson’s disease, for which antibody-based approaches such as prasinezumab are in clinical development [61, 62], and GPR52, an emerging target for Huntington’s disease [35]. GPR52 is highly expressed in the striatum and promotes stabilisation of mutant huntingtin; accordingly, genetic or pharmacological inhibition of GPR52 lowers mutant HTT levels and ameliorates Huntington’s disease-related phenotypes in preclinical models [33, 63]. GPR52 is also of broader therapeutic interest, as the oral agonist HTL0048149 (HTL’149) has recently entered clinical testing for schizophrenia [64], underscoring the translational relevance of this rhythmic striatal receptor. These findings raise the possibility that the effectiveness of neurodegenerative and psychiatric therapies may depend on the time of administration relative to target rhythmicity, opening new opportunities for chronotherapy and temporally-informed target engagement.

This work has several limitations. This is a cross-sectional atlas derived from post-mortem bulk RNA-seq data rather than a direct within-person time series, so not every oscillation can be assumed to reflect cell-autonomous circadian control. Local tissue phase may also contain non-circadian 24-hour rhythmic components from external factors such as feeding, sleep and exercise alongside canonical clock output. Thus, we do not interpret them as purely circadian, but as nycthemeral, oscillating across the 24-hour day, and comprised of circadian, entrainment and evoked artefacts. The use of bulk tissue also mixes oscillations in gene regulation with time-varying cell-state and cell-composition effects, a limitation underscored by recent single-cell studies of the human brain, which show differences across cell-types within a tissue [65]. Finally, our study measures transcript abundance only, and transcript rhythmicity does not necessarily translate into corresponding protein-level rhythmicity, because circadian regulation is extensively shaped by post-transcriptional, translational and protein-stability mechanisms [66, 67]. Although previous work suggests substantial overlap between rhythmic mRNAs and proteins [68], the extent to which these transcript-level rhythms are reflected in protein timing and abundance requires experimental validation.

Together, our results establish tissue-resolved phase estimation as a powerful framework for reconstructing human temporal biology from population-scale transcriptomic data. They reveal that local rhythmic organisation is a fundamental property of human tissues, with important implications for disease biology, target discovery and the development of chronobiology-informed therapies.

## Methods

Gene-level RNA-seq counts and sample annotations were obtained from GTEx v11. Only standard RNA-seq samples and only solid tissues were retained. Sample quality-control filters were RNA integrity number >6, total reads >40 million, uniquely mapped reads >0.8 and ischaemic time >0. Only tissues represented by at least 100 samples after filtering were retained. Within each tissue, lowly expressed genes were removed (mean raw count ≤10 across samples in that tissue), followed by normalisation and log2 transformation using edgeR [69]. Genes were further filtered by mean log2 counts per million >3 for phase inference and >0 for downstream rhythmicity analyses. Expression values were residualised using gene-wise linear models adjusting for ischaemic time, sex and Hardy death type; in sex-specific tissues, the sex variable was omitted. Our final analytic sample included 14,886 samples from 45 tissues from 927 donors.

Local and global phases were inferred from the residualised expression matrices using CHIRAL [32]. Briefly, phase inference utilised a compact set of 12 canonical clock reference genes: *DBP, PER3, TEF, NR1D2, PER1, PER2, NPAS2, ARNTL, NR1D1, CRY1, CRY2* and *CIART* as per Talamanca, Gobet [32]. CHIRAL infers circadian phase with an arbitrary phase origin and direction, thus tissue-specific phase estimates were subsequently harmonised across tissues by aligning them to the temporal ordering of core clock genes in benchmarked time-labelled human skeletal muscle data [70] and the common phase origin rotated such that *PER3* peak expression corresponded to 09:00 hours. Donor-level GPs were derived from the donor × tissue LP matrix, following the approach of Talamanca, Gobet [32]. Briefly, the GP was estimated from the dominant cluster of tissue-specific phase vectors, with agreement defined as tissue-specific phases differing by no more than 1.7 h. The GP was defined as the circular mean of the largest cluster of closely aligned tissue phases.

The residualised gene expression values were used for rhythmicity testing. Rhythmicity was assessed gene by gene using 24-hour harmonic regression, comparing an intercept-only model with a model containing sine and cosine terms: y_gi_ = α_g_ + β_g,1_cos(θ_i_) + β_g,2_sin(θi) + ε_gi_, where y_gi_ is the residualised expression value for gene *g* in sample *i*, and θ_i_ is the inferred phase assigned to that sample, derived from tissue-level LPs. The null model was y_gi_ = α_g_ + ε_gi_. Expression outliers deviating by more than 5 units from the gene-wise mean were masked before model fitting. Rhythmicity was assessed by testing whether inclusion of the sine and cosine terms significantly improved model fit relative to the intercept-only model. For each gene, phase was calculated from the arctangent of the fitted sine and cosine coefficients, amplitude as twice the square root of the sum of their squared coefficients, model *R*^2^ as the coefficient of determination of the harmonic regression model, and the Bonferroni-adjusted *P* value by multiplying the likelihood-ratio test *P* value by the number of genes tested in that tissue.

Pathway enrichment analysis was performed using Enrichr [71] on Bonferroni-significant rhythmic gene sets for each tissue. Functional enrichment was evaluated against KEGG 2026 [72], drug-gene interaction enrichment against DGIdb 2022 [73], and transcription factor enrichment against the ChEA database [74], with significance defined as FDR adjusted *P* < 0.05.

